# Highly pathogenic avian influenza A(H5N1) virus clade 2.3.4.4b in domestic ducks, Indonesia, 2022

**DOI:** 10.1101/2023.07.10.548369

**Authors:** Hendra Wibawa, Putut Eko Wibowo, Arif Supriyadi, Lestari, Jessiaman Silaban, Aziz Ahmad Fuadi, Anna Januar Fiqri, Retno Wulan Handayani, Sri Handayani Irianingsih, Zaza Fahmia, Herdiyanto Mulyawan, Syafrison Idris, Nuryani Zainuddin

**Affiliations:** Disease Investigation Center Wates, Yogyakarta, Indonesia; Disease Investigation Center Banjarbaru, Banjarbaru, South Kalimantan, Indonesia; Directorate General of Livestock and Animal Health Services, Jakarta, Indonesia

**Keywords:** HPAI H5N1, clade 2.3.4.4b, ducks, whole genome sequencing, Indonesia

## Abstract

Highly pathogenic avian influenza A(H5N1) clade 2.3.4.4b viruses were detected and isolated from domestic ducks in South Kalimantan, Indonesia during April 2022. The viruses were genetically similar to clade 2.3.4.4b H5N1 viruses recently detected in East Asia from 2021-2022. Further investigation is necessary to determine the source of virus incursion.

---

H5N1 subtype of avian influenza (AI) virus A/goose/Guangdong/1/96 (Gs/GD/96) lineages have caused highly pathogenic AI (HPAI) outbreaks in poultry since 1996. In 2008, various novel reassortant viruses bearing the genetic backbone of the hemaggutinin (HA) of Gs/GSD/96 from clade 2.3.4 with different combination of neuramindase (NA) were identified in domestic duck and live poultry markets in China including H5N2, H5N5, H5N6, H5N8 (Lee et al., 2017). Clade 2.3.4 continued to evolve resulting several fourth order lineages, including clade 2.3.4.4, which have undergone genetic reassortment with other H5N1 clades and low pathogenic AI (LPAI) viruses resulting eight genetic groups (2.3.4.4a-h) and multiple genotypes (Lee et al., 2017). Clade 2.3.4.4 of H5N8 subtype viruses have spread across many countries from Asia to Europe, Africa, and North America (Lee et al, 2017; Kwon, et al., 2023) and repeated outbreaks due to infection of H5N8 clade 2.3.4.4 group B (2.3.4.4b) viruses have been reported between 2016 and mid-2020 (Yehia et al, 2018; Engelsma et al, 2022). However, a new strain of HPAI H5N1 clade 2.3.4.4b virus emerged in late 2020 which led to an increase the scale of wild bird outbreaks and poultry worldwide and this virus has almost entirely replaced H5N8 clade 2.3.4.4b globally since late 2021 (Venkatesan, 2023; Wiley and Barr, 2022)

In April 2022, several poultry cases were reported from small-holder duck farms located next to marshes in Hulu Sungai Utara District, South Kalimantan Province, Indonesia (Appendix, Figure 1). We collected 18 oropharyngeal swabs which pooled in four viral transport media and two tissue samples from dead birds from these farms. All samples were detected AI positive by PCR, but only 3 pooled swabs could be isolated from embryonic chicken eggs. They were then characterized using Illumina sequencing platform followed by phylogenetic and sequence analysis based on complete genome sequencing. We conducted the hemagglutination inhibition (HI) assay for the virus isolates and performed necropsy and hematoxylin and eosin staining for gross- and histo-pathology of collected tissues from dead ducks.

Whole genome sequence of three virus isolates, A/duck/Hulu Sungai Utara/A0522064-06/2022, A/duck/Hulu Sungai Utara/A0522064-03-04/2022 and A/duck/Hulu Sungai Utara/A0522067-06-07/2022, have been deposited to GISAID under ID isolate numbers EPI_ISL_17371282, EPI_ISL_17371283, and EPI_ISL_17371284, respectively. Phylogenetic analysis of the HA gene showed that all three viruses clustered with recent H5 clade 2.3.3.4b viruses from Asia and Europe. However, they appeared to be more genetically close related to H5N1 clade 2.3.4.4b viruses from wild birds and poultry in China, South Korea, and Japan between October 2021 and February 2022 (Figure). The tree topology of the other gene segments (PB2, PB1, PA, NP, NA, MP, and NS) showed that all three viruses were also situated close to H5N1 clade 2.3.4.4b from China, South Korea, and Japan (Appendix Figures 2-5). These virus isolates shared 99.8-100% nucleotide sequences similarity in each viral segment and all were identified as HPAI on the basis of multiple basic amino acid sequences detected in the HA cleavage site (REKRRKR|G). None of these have molecular determinants related with increase binding affinity or efficient replication in mammals including humans (Chutinimitkul et al., 2010; Suttie et al., 2019) (Appendix Table 1). The BLAST tool search (https://www.ncbi.nlm.nih.gov/blast) and pairwise distance analysis showed that all eight gene segments had 98.4-99.8% nucleic acid sequence identities to those of the H5N1 clade 2.3.4.4b viruses from China, South Korea, and Japan (Table) indicating that they had a close common ancestor.

The HI assay revealed low reactivities of the virus isolates against representative antisera from HPAI viruses that have been circulating in poultry in Indonesia, including the H5N1 vaccines that are currently used (Appendix, Table 2). The gross- and histo-pathology in naturally infected ducks showed multiorgan haemorrhages with prominent lesion in tissues were congestion and necrosis in parenchymal cells often accompanied with inflammatory cell infiltrates (Appendix, Figure 6)

Wild migratory birds have been considered play a role in the intercontinental spread of HPAI H5Nx clade 2.3.4.4 (Lee et al, 2017; Caliendo et al, 2022, Zhang et al, 2023). This study reports the identification of HPAI H5N1 clade 2.3.4.4b viruses isolated from domestic ducks in South Kalimantan, Indonesia. Although we could not yet determine the source of virus incursion to this area, the role of wild aquatic birds on virus transmission cannot be rule out since the infected farms were connected to water course from the marshes which provide opportunity for intermingle contacts between naturally infected wild birds to naïve ducks or these ducks could be infected through contaminated water.

Limitation of data on AI virus genome sequence and wild bird surveillance hampering us to determine the exact role of wild birds in the spread of HPAI H5N1 clade 2.3.4.4b into Indonesia. This highlights the importance of both epidemiology and molecular surveillance on these species to support better preparedness for such pandemic threats due to continued AI virus evolution.

## Supporting information

Technical Appendix

## Acknowledgments

We are grateful to authors who deposited sequences in GISAID’s EpiFlu Database (http://www.gisaid.org, Appendix Table 3. Sequence Acknowledgements Table). We thank to DIC Banjabaru’s staff for conducting field investigation and initial laboratory diagnostics and also DIC Wates’s staff for virus isolation, serology testing and whole genome sequencing. This study was supported by the Directorate General of Livestock and Animal Health Services (DGLAHS) of Ministry of Agriculture, Indonesia and was also supported for sequencing reagents by the United Nations Food and Agriculture Organization Emergency Centre for Transboundary Animal Diseases, Jakarta, Indonesia.

## About Author

Dr Wibawa is a veterinarian at the Disease Investigation Centre Wates, DGLAHS Ministry of Agriculture, Indonesia. His research interests are molecular diagnostics and epidemiology of influenza viruses in animals.

## Figure Legend

Phylogenetic tree of the HA gene of H5N1 clade 2.3.4.4b viruses isolated from domestic ducks (indicated in red taxa) during poultry outbreaks in South Kalimantan, Indonesia in April 2022. The evolutionary history was inferred by using Maximum Likelihood method using the best fit substitution model (GTR+G) involving 67 HA-H5 sequences from GISAID (http://www.gisaid.org) with a total of 1656 positions in the final dataset. The tree is drawn to scale, with branch lengths measured in the number of substitutions per site (0.01) shown in bottom left.

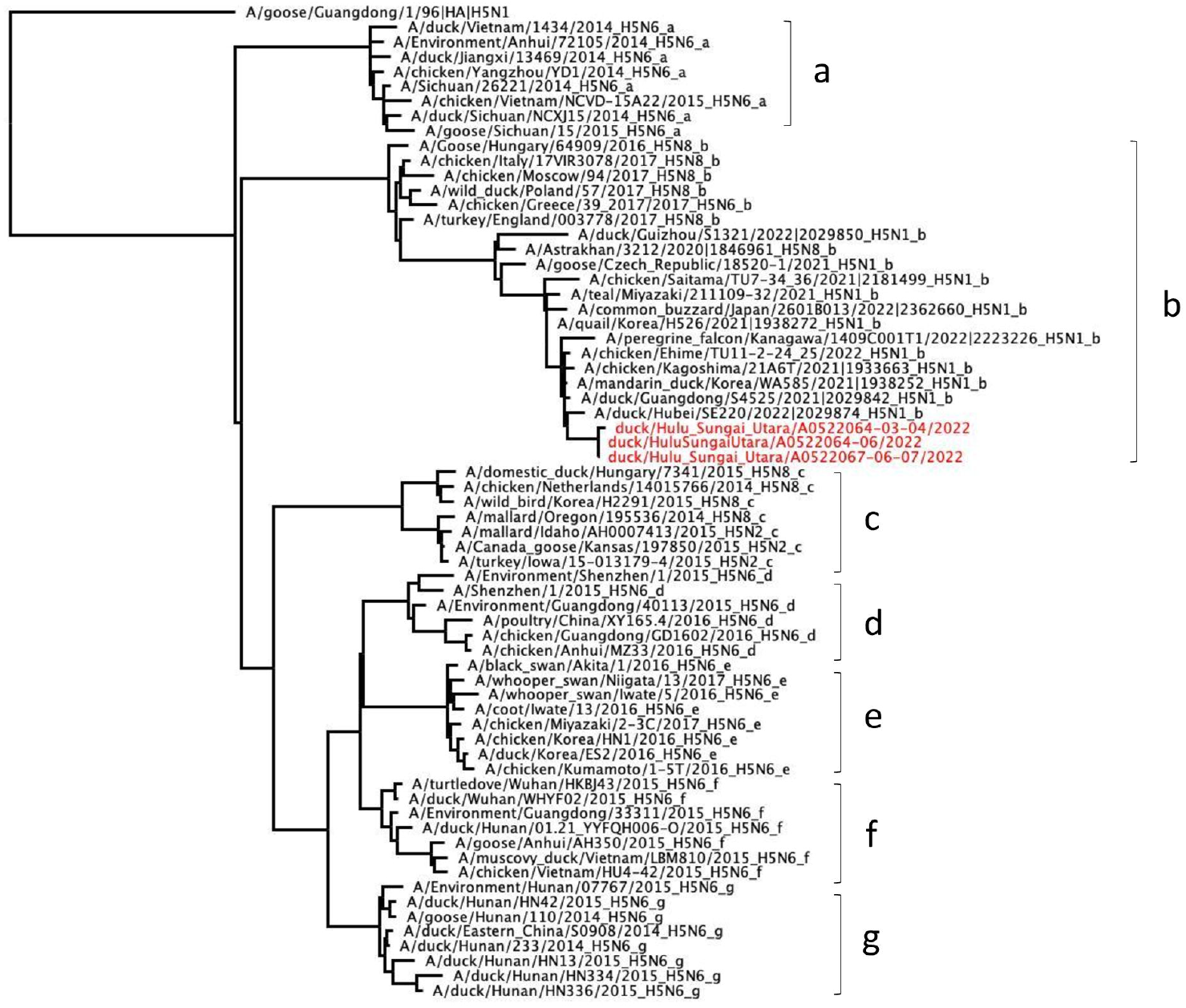

## Table Title

Sequence homology of each gene segment of the virus isolates to other H5N1 clade 2.3.4.4b viruses from China, Korea, and Japan, October 2021 to February 2022.

| Virus Name<br>(GISAID ID No. \| Collection Date) | Nucleic acid similarity (%) |  |  |  |  |  |  |  |
| --- | --- | --- | --- | --- | --- | --- | --- | --- |
|  | PB2 | PB1 | PA | HA | NP | NA | MP | NS |
| Between the virus isolates<br>(EPI_ISL_17371282, EPI_ISL_17371283,<br>EPI_ISL_17371284 \| 2022-04-05) | 100 | 99.8-99.9 | 99.8-99.9 | 99.9-100 | 100 | 100 | 100 | 99.8-100 |
| A/mandarin duck/Korea/WA585/2021<br>(EPI_ISL_6959592 \| 2021-10-26) | 99.6 | 99.5-99.6 | 99.6-99.7 | 99.2-99.3 | 99.6 | 99.8 | 99.6 | 99.4 |
| A/quail/Korea/H526/2021<br>(EPI_ISL_6959593 \| 2021-11-08) | 99.4 | 99.2 | 99.2-99.3 | 99.0-99.1 | 99.3 | 99.6 | 99.3 | 99.2 |
| A/duck/Guangdong/S4525/2021<br>(EPI_ISL_12572655 \| 2021-12-08) | 99.6 | 99.6 | 99.3-99.4 | 99.2-99.3 | 99.5 | 99.7 | 99.7 | 99.2 |
| A/duck/Hubei/SE220/2022<br>(EPI_ISL_12572659 \| 2022-01-10) | 99.6 | 99.5 | 99.3-99.4 | 99.0-99.2 | 99.5 | 99.5 | 99.6 | 99.2 |
| A/duck/Guizhou/S1321/2022<br>(EPI_ISL_12572656 \| 2022-02-22) | 99.6 | 99.6 | 99.5-99.6 | 97.2-97.3 | 99.5 | 99.4 | 99.8 | 99.2 |
| A/chicken/Kagoshima/21A6T/2021<br>(EPI_ISL_6829533 \| 2021-11-12) | 99.6 | 99.6 | 99.6-99.7 | 99.1-99.2 | 99.6 | 99.7 | 99.8 | 99.4 |
| A/chicken/Saitama/TU7-34,36/2021<br>(EPI_ISL_15063425 \| 2021-12-07) | 99.6 | 99.3-99.4 | 99.1-99.2 | 98.4-98.6 | 99.1 | 99.6 | 99.3 | 99.0 |
| A/teal/Miyazaki/211109-32/2021<br>(EPI_ISL_15613494 \| 2021-11-09) | 99.4 | 99.3 | 99.3 | 98.7-98.9 | 99.1 | 99.6 | 99.2 | 99.2 |
| A/chicken/Ehime/TU11-2-24 25/2022<br>(EPI_ISL_15063431 \| 2022-01-04) | 99.8 | 92.5 | 99.5-99.6 | 99.2-99.3 | 99.3 | 99.6 | 99.7 | 99.3 |
| A/common buzzard/Japan/2601B013/2022<br>(EPI_ISL_16831015 \| 2022-01-27) | 99.6 | 99.3 | 99.3-99.4 | 98.6-98.7 | 99.3 | 99.6 | 99.3 | 99.2 |

