## Supplementary material for "Highly pathogenic avian influenza A(H5N1) virus clade 2.3.4.4b in domestic ducks, Indonesia, 2022": Technical Appendix

**Material and Methods**

**Samples**

Eighteen oropharyngeal swabs were collected and pooled in four viral transport media and two tissue samples from dead and sick domestic ducks in 4^th^ of June 2022 from three small-holder duck farms which connected to marshes in Hulu Sungai Utara District in South Kalimantan Province of Indonesia by Disease Investigation Center (DIC) Banjarbaru (the location and the picture of duck farm displayed in Appendix Figure 1). All original pooled swab samples detected positive for influenza A virus were sent to the National Avian Influenza Reference Laboratory at Disease Investigation Center (DIC) Wates for further virus characterizations. Samples from three pooled swabs that could be isolated from 9- to 10-day-old specific pathogen-free chicken embryonic eggs (A/duck/Hulu Sungai Utara/A0522064-06/2022, A/duck/Hulu Sungai Utara/A0522064-03-04/2022, and A/duck/Hulu Sungai Utara/A0522067-06-07/2022) were then characterized antigenically by hemagglutination inhibition (HI) assay and genetically by whole genome sequencing for avian influenza virus.

**Genome Sequencing and Phylogenetic Analysis**

We performed RNA extraction using QIAamp Viral RNA Mini Kit, followed by the multisegment reverse-transcription polymerase chain reaction with specific primers MBTuni-12 and MBTuni-13 to amplify all eight gene segments of avian influenza virus (Zhou et al., 2009). The DNA libraries were prepared by using Nextera-XT DNA Library Preparation Kit according to manufacturer`s instructions (Illumina). The whole-genome sequencing was carried out under MiSeq next-generation sequencing (NGS) machine with MiSeq Reagent Kit v3 (Illumina). Validation and assembly of NGS nucleotide sequences were performed by using Geneious Prime v2022.2.1 (Geneious).

Complete genome sequencing of A/duck/Hulu Sungai Utara/A0522064-06/2022, A/duck/Hulu Sungai Utara/A0522064-03-04/2022, and A/duck/Hulu Sungai Utara/A0522067-06-07/2022 have been deposited in GISAID under Isolate ID accession numbers EPI_ISL_17371282, EPI_ISL_17371283, and EPI_ISL_17371284, respectively. The PhyML maximum-likelihood phylogenetic amalysis for each of the gene segments were reconstructed using the general time-reversible nucleotide substitution model in Unipro UGENE v.46 (<https://ugene.net/>) to produce newick trees. Final dendrograms were generated and visualized in FigTree v.1.4.4 (<https://github.com/rambaut/figtree/releases>). We performed initial the BLAST tool search (<https://www.ncbi.nlm.nih.gov/blast>) followed by nucleotide identity analysis which calculated from the output of pairwise distance analysis of each gene segment in MEGA X (Kumar et al., 2005).

**Hemagglutination Inhibition Test**

We performed HI assays following the WOAH standard method (<https://www.woah.org/fileadmin/Home/eng/Health_standards/tahm/3.03.04_AI.pdf>) to test the reactivity these three virus isolates against some representative antisera from H5N1 clade 2.1.3.2 virus strains (very few still detected in poultry), clade 2.3.2.1c virus strains (dominant circulating virus clade in poultry) and from H5N6 clade 2.3.4.4 (A/duck/Laos/XBY004/2014).

**Gross Pathology and Histopathology**

Carcasses from the dead ducks were necropsied by veterinary pathologists within the post-mortem facility in DIC Banjarbaru. Tissue samples included brain, lungs, heart, liver, spleen, pancreas, intestines, and kidney were processed and embedded into wax by routine histological processes. One section of each tract was cut by standard microtomy methods, and consecutive 4-mm-thick sections were stained with hematoxylin and eosin (HE).

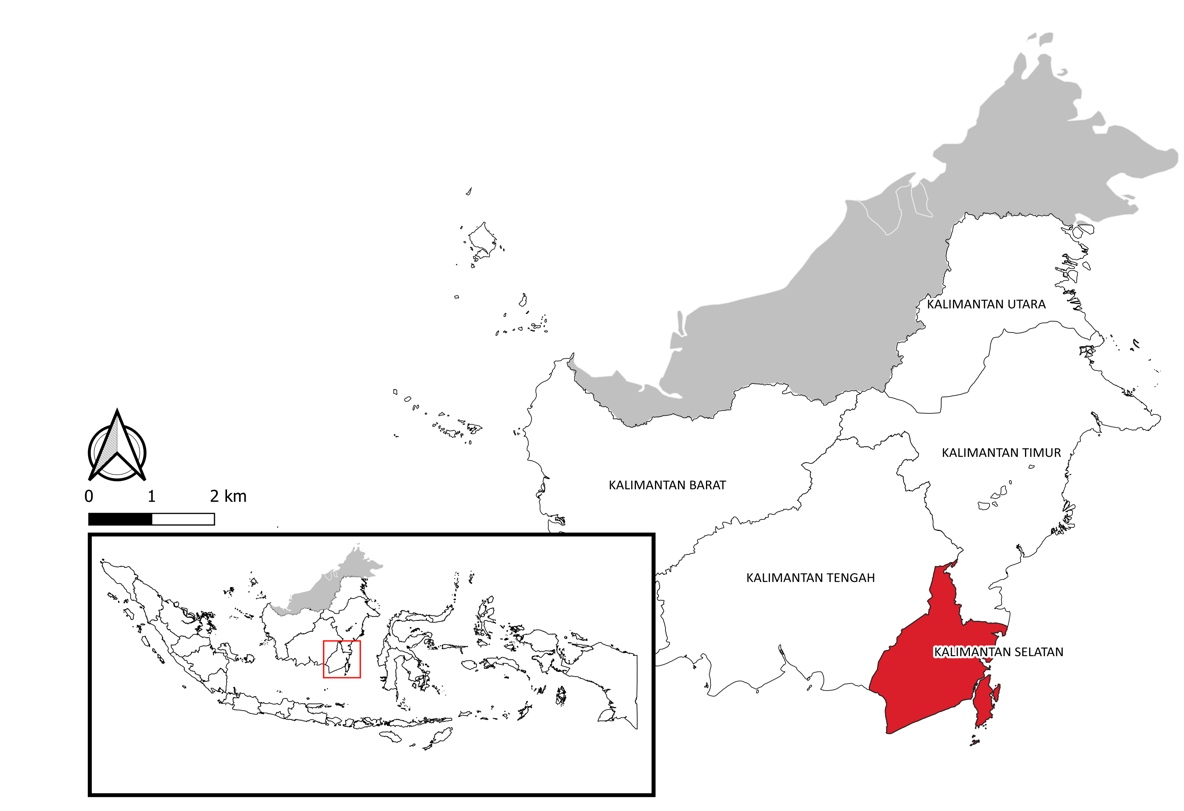

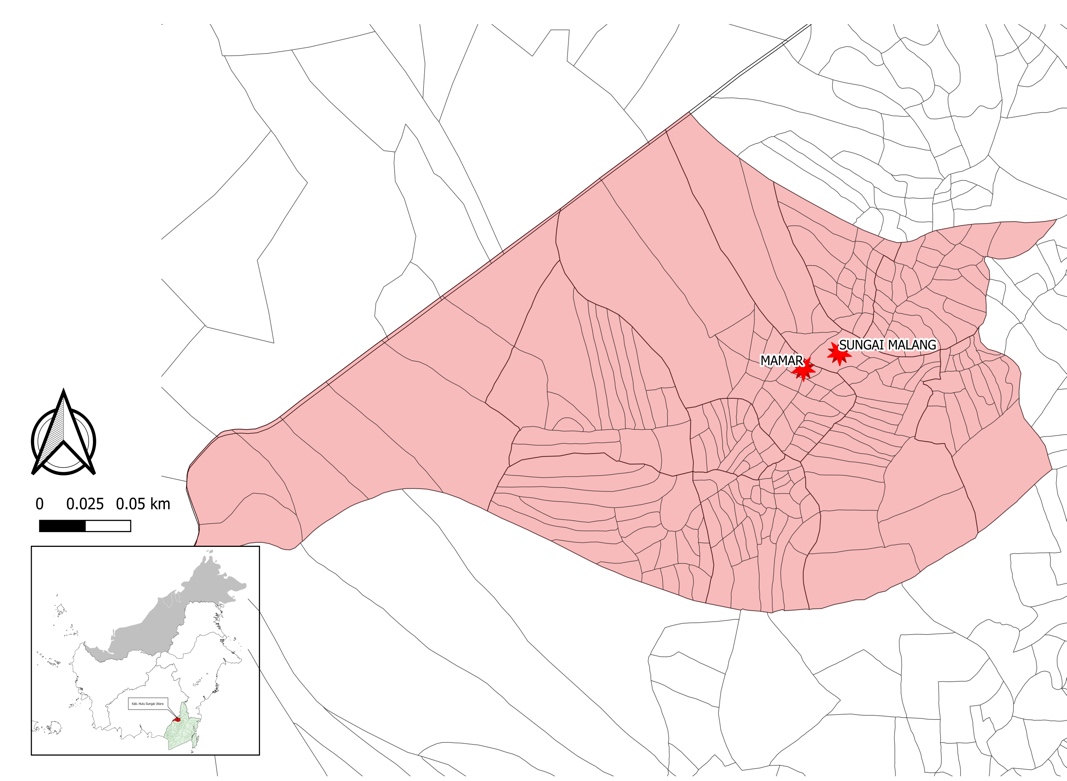

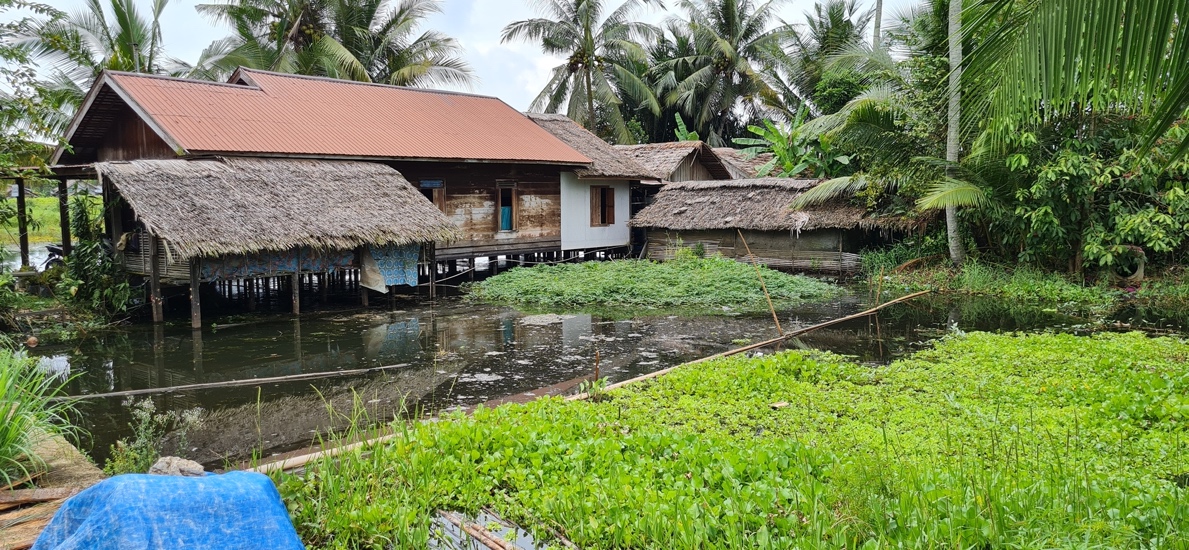

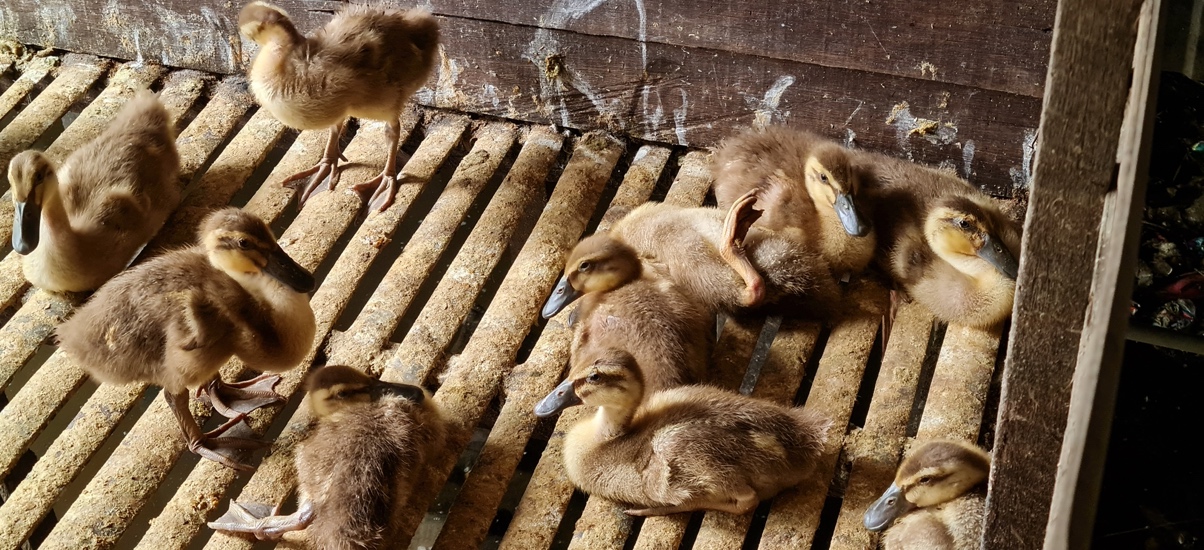

**A**

**B**

**C**

**D**

Appendix Figure 1. Location of poultry cases in small-holder duck farms in South Kalimantan Province, Indonesia (A). Map showing the exact places where the samples were collected form infected duck farms in Hulu Sungai Utara District (B). A photo showing farms located above a water flow from marshes (C). High mortality found up to 60% particularly in young ducks, often with neurological signs such as torticollis and paralysis (D).

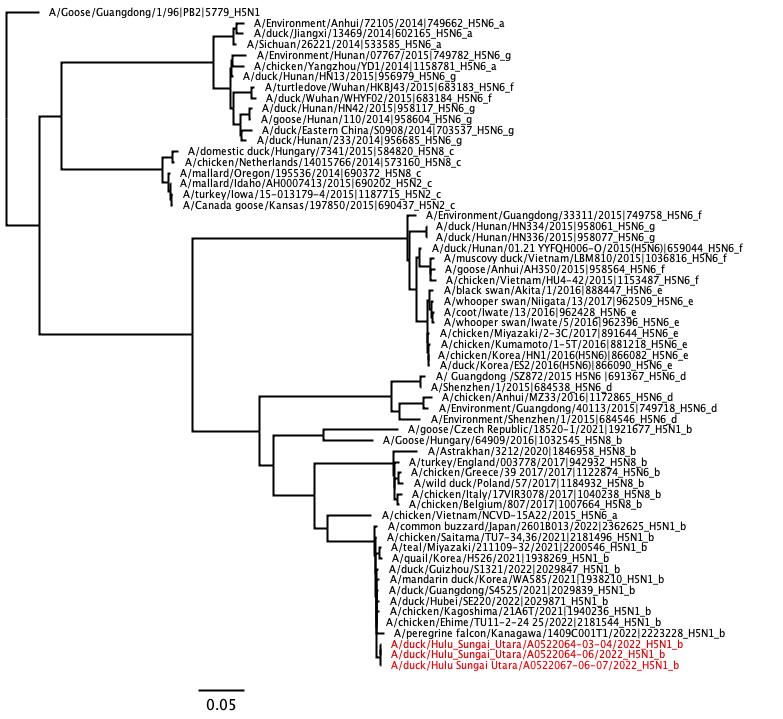

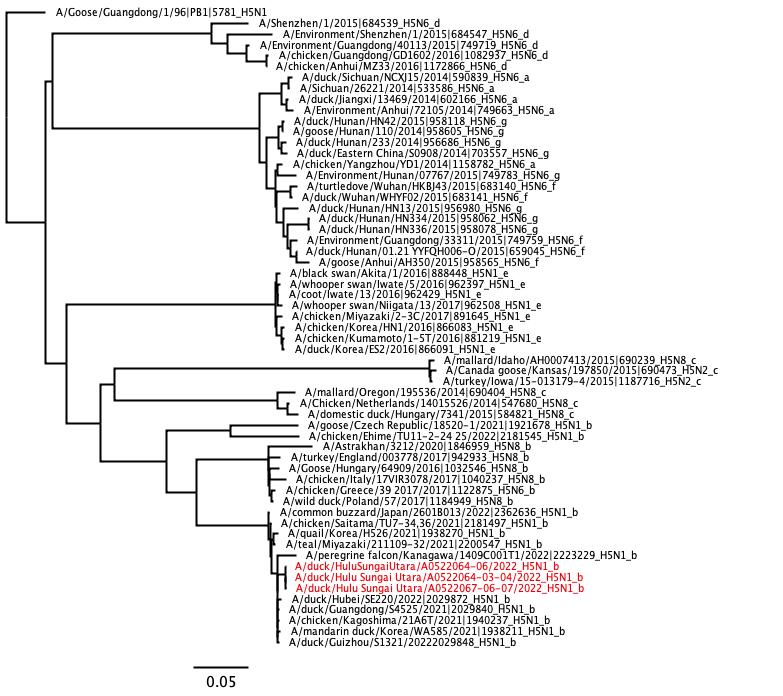

**PB2**

**PB1**

Appendix Figure 2. Maximum likelihood tree of PB2 and PB1 gene segments of clade 2.3.4.4 H5 subtypes including clade 2.3.4.4b H5N1 isolated from domestic ducks (indicated in red taxa) during poultry outbreaks in South Kalimantan, Indonesia in April 2022. The tree is drawn to scale, with branch lengths measured in the number of substitutions per site shown in bottom left.

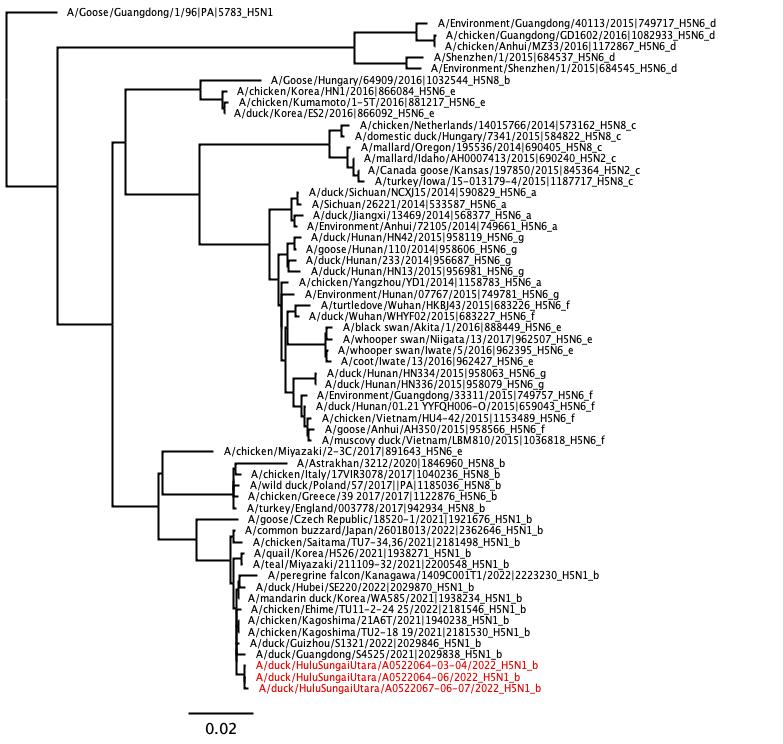

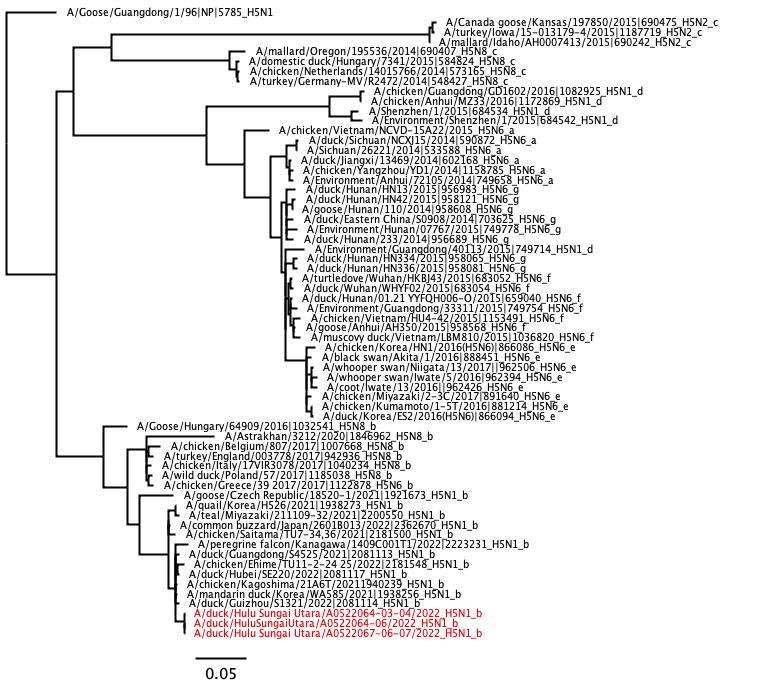

**PA**

**NP**

Appendix Figure 3. Maximum likelihood tree of PA and NP gene segments of clade 2.3.4.4 H5 subtypes including clade 2.3.4.4b H5N1 isolated from domestic ducks (indicated in red taxa) during poultry outbreaks in South Kalimantan, Indonesia in April 2022. The tree is drawn to scale, with branch lengths measured in the number of substitutions per site shown in bottom left.

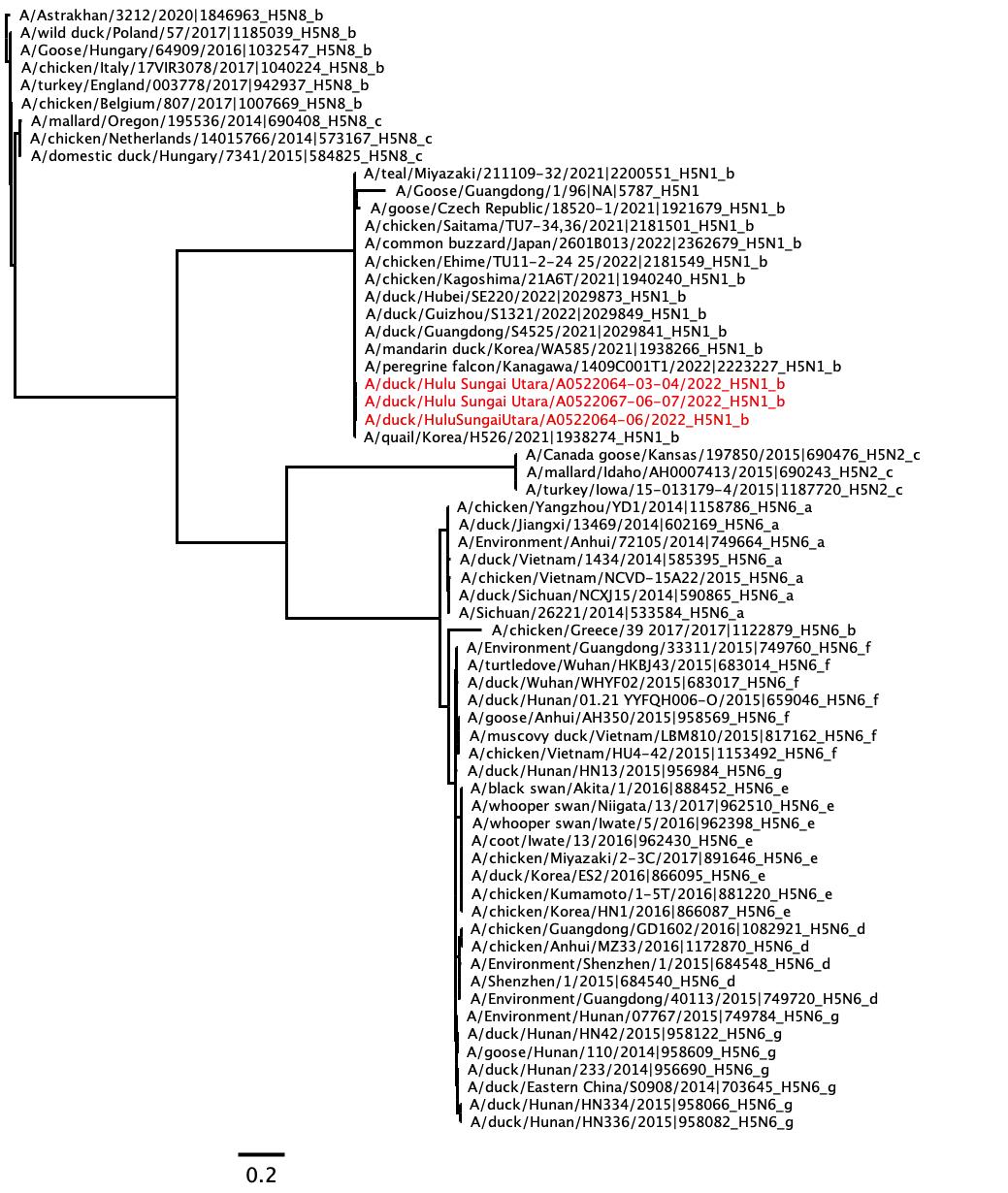

**N8**

**N1**

**N2**

**N6**

Appendix Figure 4. Maximum likelihood tree of NA gene segments of clade 2.3.4.4 H5Nx subtypes including clade 2.3.4.4b H5N1 isolated from domestic ducks (indicated in red taxa) during poultry outbreaks in South Kalimantan, Indonesia in April 2022. The tree is drawn to scale, with branch lengths measured in the number of substitutions per site shown in bottom left. Nx: N1, N2, N6, N8.

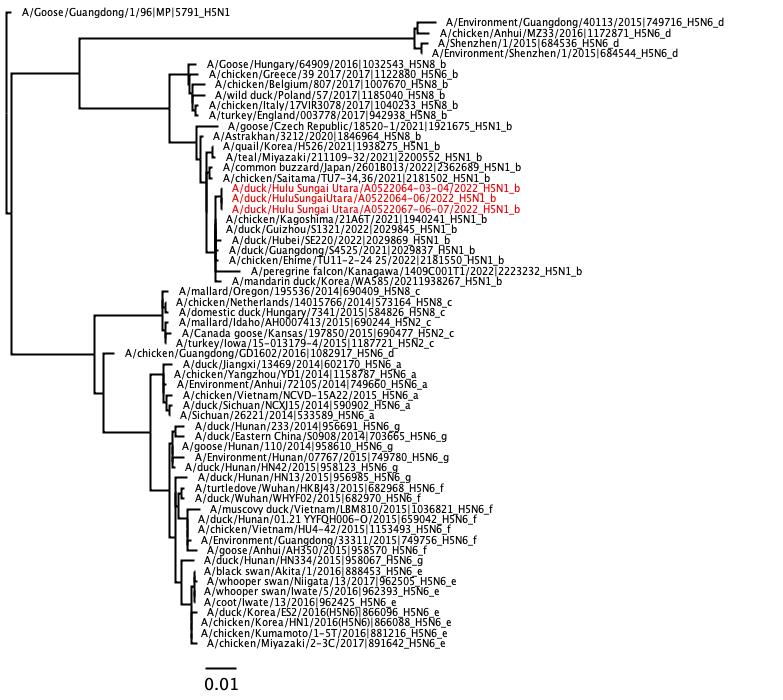

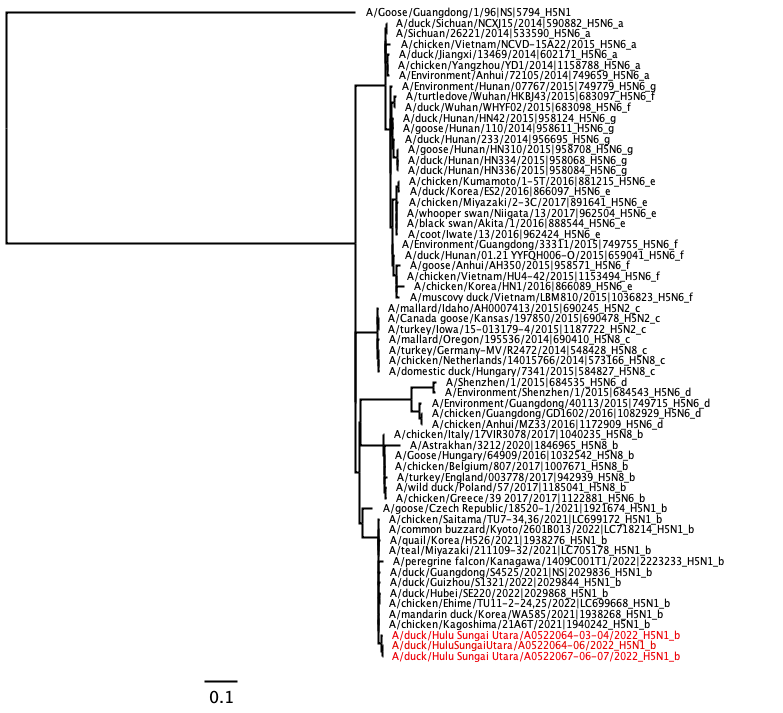

**MP**

**NS**

Appendix Figure 5. Maximum likelihood tree of MP and NS gene segments of clade 2.3.4.4 H5 subtypes including clade 2.3.4.4b H5N1 isolated from domestic ducks (indicated in red taxa) during poultry outbreaks in South Kalimantan, Indonesia in April 2022. The tree is drawn to scale, with branch lengths measured in the number of substitutions per site shown in bottom left.

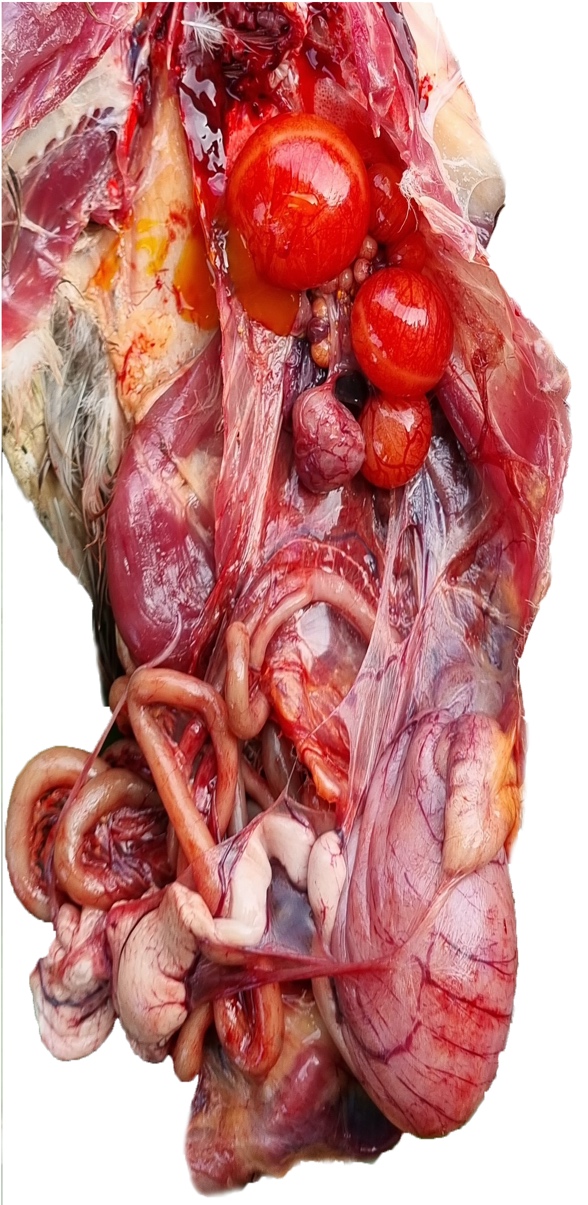

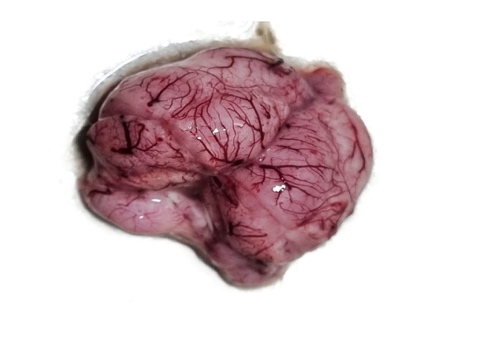

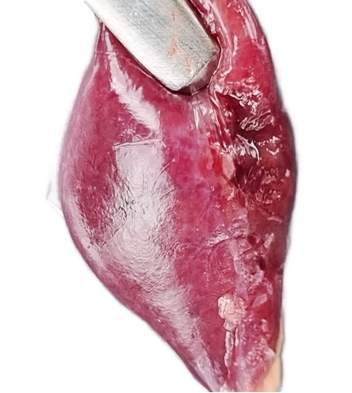

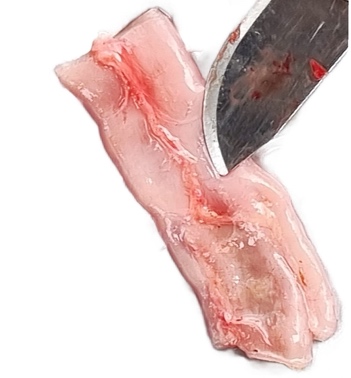

**(A)A**

**(B)**

**(C)**

**(D)**

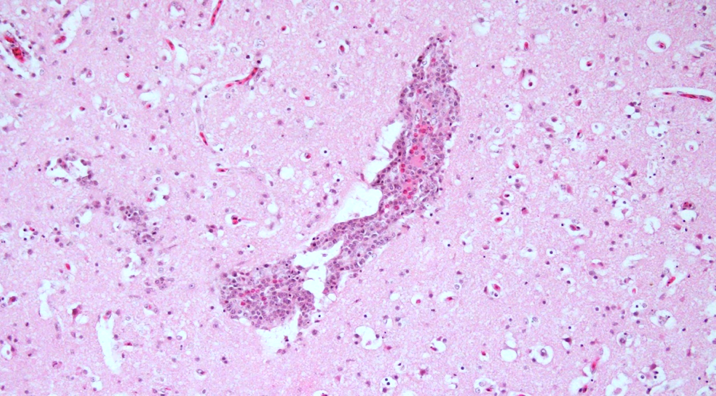

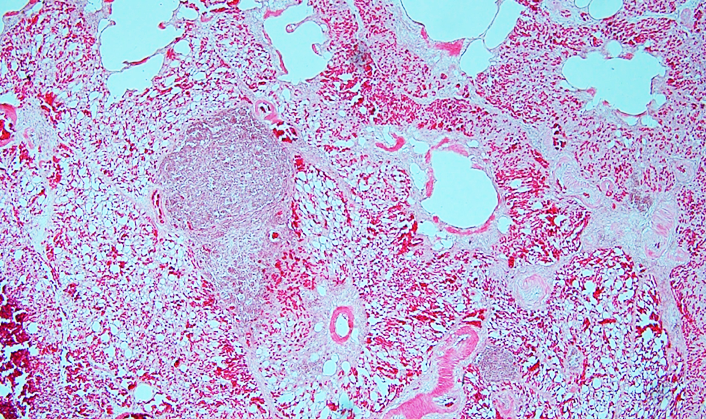

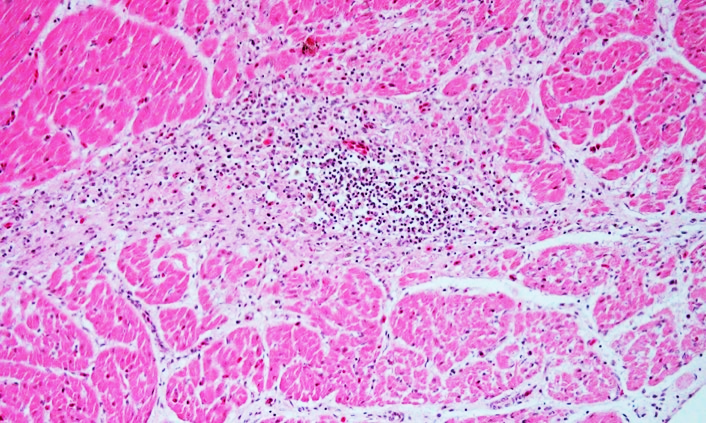

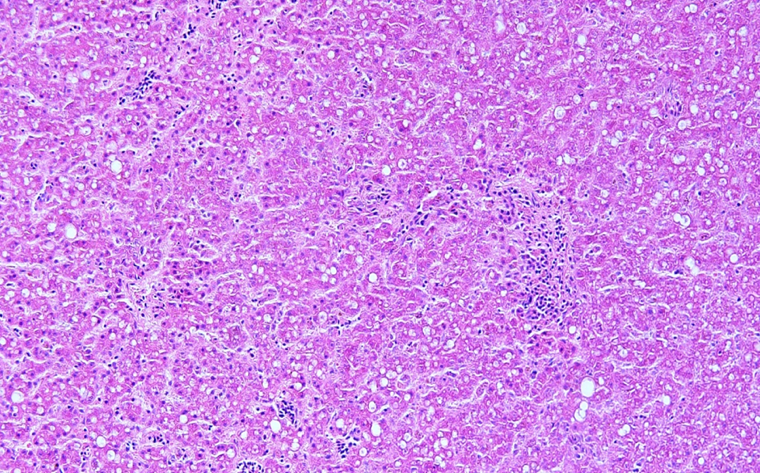

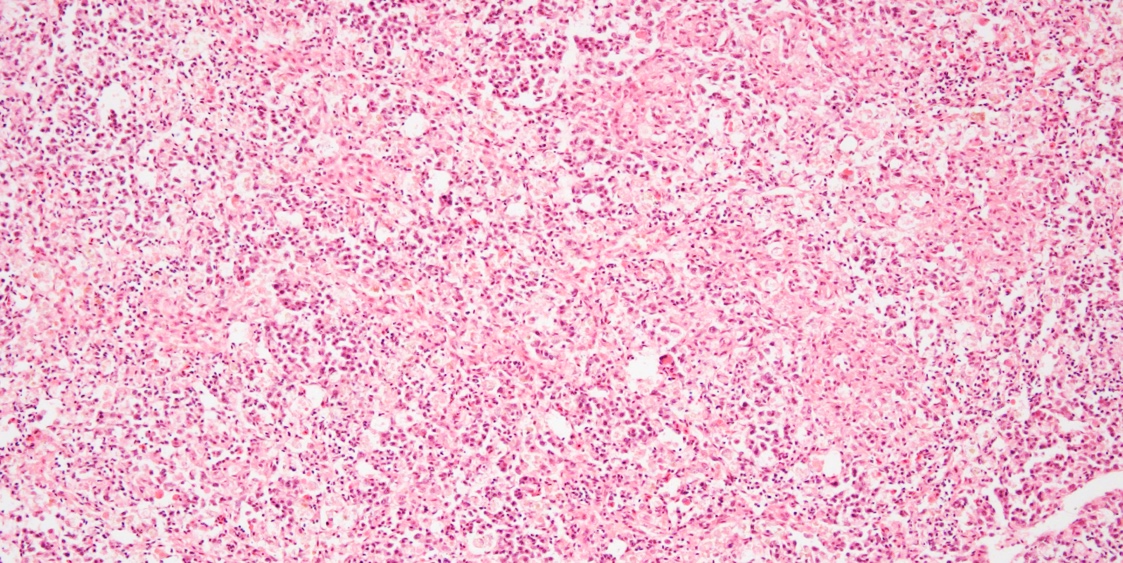

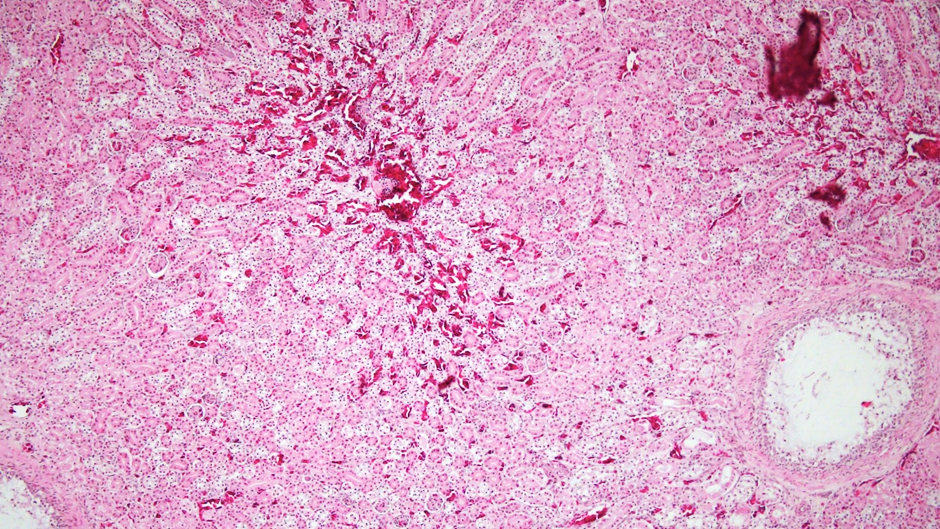

**(E)**

**(F)**

**(G)**

**(H)**

**(I)**

**(J)**

Appendix Figure 6. Gross pathology (A-D) and histopathology (E-J) of ducks naturally infected with HPAI H5N1 clade 2.3.4.4b in South Kalimantan, Indonesia in April 2022. Haemorrhages and acute necrosis observed in visceral organs (A) including in brain (B), spleen (C), intestine (D). Histopathology of brain shows congestion, edema, gliosis, and mononuclear cell perivascular cuff (E), lungs show congestion, perivascular edema, and alveolar lumen constriction (F), hearth shows focal necrosis and inflammatory cell infiltrates (G), liver shows focal necrosis, inflammatory cell infiltrates, and lipid degeneratopn (H), pancreas shows mild haemorrhages and focal necrosis (I), kidney shows interstitial haemorrhages, congestion, and focal necrosis (J).

Appendix Table 1. Amino acid changes related an increase binding activity and replication in mammalian cells or increase virulence in mammals

| **Virus Name** | **PB2** | | | **PB1** | **PB1-**  **F2** | **PA** | **HA** | | | | | **NA** | **M2** | **NS** | | |
| --- | --- | --- | --- | --- | --- | --- | --- | --- | --- | --- | --- | --- | --- | --- | --- | --- |
|  | **Q59K** | **E627K** | **D701N** | **N105S** | **N66S** | **T91I** | **Cleavage**  **site** | **Q192R** | **Q222L** | **S223N** | **G224S** | **Stalk**  **Del.** | **S31N** | **NS Del. 80 -84** | **P42S** | **PDZ**  **motif** |
| A/Viet Nam/1203/2004 (Clade 3A.1) | Q | **K** | D | N | N | T | RERRRKK-R\|G | Q | Q | S | G | **Yes** | **N** | **Yes** | **S** | ESEV |
| A/Jiangsu/NJ210/2023 (Clade 2.3.3.4b) | Q | E | D | N | N | T | REKRRK-R\|G | **K** | Q | **R** | G | No | S | No | **S** | ESEV |
| A/duck/Hulu Sungai Utara/A0522064-06/2022 | Q | E | D | N | N | T | REKRRK-R\|G | **K** | Q | **R** | G | No | S | No | **S** | ESEV |
| A/duck/Hulu Sungai Utara/A0522064-03-04/2022 | Q | E | D | N | N | T | REKRRK-R\|G | **K** | Q | **R** | G | No | S | No | **S** | ESEV |
| A/duck/Hulu Sungai Utara/A0522067-06-07/2022 | Q | E | D | N | N | T | REKRRK-R\|G | **K** | Q | **R** | G | No | S | No | **S** | ESEV |

Appendix Table 2. Hemagglutinin inhibition (HI) assay results using two fold serial dilution of antisera using WOAH/OIE Manual for avian influenza including HPAI (<https://www.woah.org/fileadmin/Home/eng/Health_standards/tahm/3.03.04_AI.pdf>)

| Antisera | Isolate Antigens | | |
| --- | --- | --- | --- |
|  | A/duck/Hulu Sungai Utara/A0522064-06/2022 | A/duck/Hulu Sungai Utara/A0522064-03-04/2022 | A/duck/Hulu Sungai Utara/A0522067-06-07/2022 |
| A/chicken/West Java/PWT-WIJ/2006*  (H5N1 Clade 2.1.3.2 EPI_ISL_12700530\|) | < 4 | < 4 | < 4 |
| A/chicken/Barru/BBVM 41-13/2013 (H5N1 clade 2.1.3.2a\| EPI_ISL_17767706) | 16 | 16 | 16 |
| A/duck/Sukoharjo/BBVW-1428-9/2012** (H5N1 clade 2.3.2.1c\| EPI_ISL_266808) | 16 | 32 | 32 |
| A/chicken/Tanggamus/031711076 - 65/2017** (H5N1 clade 2.3.2.1c\| EPI_ISL_17767763) | 16 | 32 | 16 |
| A/duck/Laos/XBY004/2014  (H5N6 clade 2.3.4.4\| EPI_ISL_168385) | 8 | 16 | 8 |

* One of HPAI H5N1 clade 2.1.3.2 vaccine seed strains used in Indonesia from 2009 to date (A/chicken/West Java/PWT-WIJ/2006)

** HPAI H5N1 clade 2.3.2.1c vaccine seed strain from 2012-2020 (A/duck/Sukoharjo/BBVW-1428-9/2012) and from 2021 to date (A/chicken/Tanggamus/031711076 - 65/2017)

Appendix Table 3. Sequence acknowledgement table from GISAID

| **No** | **Isolate name** | **Isolate-ID** | **Country** | **Collection date** | **Originating Lab** | **Submitting Lab** | **Authors** |
| --- | --- | --- | --- | --- | --- | --- | --- |
| 1 | A/Goose/Guangdong/1/96 | [EPI_ISL_1254](https://platform.epicov.org/epi3/start/EPI_ISL/1254) | China | 1996-Jan-01 |  | Import from public-domain |  |
| 2 | A/duck/Vietnam/1434/2014 | [EPI_ISL_177703](https://platform.epicov.org/epi3/start/EPI_ISL/177703) | Vietnam | 2014-Nov-18 |  | Import from public-domain | Hatamachi,J.; Ogaswara,K.; Chu,D.H.; Okamatsu,M.; Sakoda,Y.; Kida,H.; Ogasawara,K. |
| 3 | A/Environment/Anhui/72105/2014 | [EPI_ISL_219801](https://platform.epicov.org/epi3/start/EPI_ISL/219801) | China | 2014-May-20 |  | WHO Chinese National Influenza Center | Wang,Dayan;Li,Xiaodan;Zou,Shumei;Zhang,Ye;Bo,Hong;Li,Xiyan;Chen,Wenbing;Yang, Lei;Shu,Yuelong |
| 4 | A/duck/Jiangxi/13469/2014 | [EPI_ISL_173478](https://platform.epicov.org/epi3/start/EPI_ISL/173478) | China | 2014-Mar-30 |  | Import from public-domain | Ma,C.; Lam,T.T.Y.; Chai,Y.; Wang,J.; Fan,X.; Hong,W.; Zhang,Y.; Li,L.; Liu,Y.; Smith,D.K.; Webby,R.J.; Peiris,J.S.M.; Zhu,H.; Guan,Y. |
| 5 | A/chicken/Yangzhou/YD1/2014 | [EPI_ISL_295144](https://platform.epicov.org/epi3/start/EPI_ISL/295144) | China | 2014-Sep-01 |  | Import from public-domain | Li,J.; Gu,M.; Sun,W.; Liu,K.; Gao,R.; Liu,D.; Hu,J.; Wang,X.; Hu,S.; Liu,X. |
| 6 | A/Sichuan/26221/2014 | [EPI_ISL_163493](https://platform.epicov.org/epi3/start/EPI_ISL/163493) | China | 2014-Apr-21 |  | WHO Chinese National Influenza Center |  |
| 7 | A/chicken/Vietnam/NCVD-15A22/2015 | [EPI_ISL_244487](https://platform.epicov.org/epi3/start/EPI_ISL/244487) | Vietnam | 2015-Apr-02 |  | Import from public-domain | Davis,T.; Jang,Y. |
| 8 | A/duck/Sichuan/NCXJ15/2014 | [EPI_ISL_179647](https://platform.epicov.org/epi3/start/EPI_ISL/179647) | China | 2014-Apr-27 |  | Import from public-domain | Bi,Y.; Chen,Q.; Chen,J. |
| 9 | A/goose/SiChuan/15/2015 | [EPI_ISL_255850](https://platform.epicov.org/epi3/start/EPI_ISL/255850) | China | 2015-Apr-07 |  | Wuhan Institute of Virology |  |
| 10 | A/Goose/Hungary/64909/2016 | [EPI_ISL_271713](https://platform.epicov.org/epi3/start/EPI_ISL/271713) | Hungary | 2016-Dec-14 | National Food Chain Safety Office Veterinary Diagnostic Directorate Laboratory for Molecular Biology | Danam.Vet.Molbiol | Adam, Dan |
| 11 | A/chicken/Italy/17VIR3078/2017 | [EPI_ISL_273846](https://platform.epicov.org/epi3/start/EPI_ISL/273846) | Italy | 2017-Apr-07 | Istituto Zooprofilattico Sperimentale Delle Venezie | Istituto Zooprofilattico Sperimentale Delle Venezie | Bianca,Zecchin; Alice,Fusaro; Gianpiero,Zamperin; Alessia,Schivo; Annalisa,Salviato; Sabrina,Marciano; Silvia,Ormelli; Calogero,Terregino; Isabella,Monne |
| 12 | A/chicken/Moscow/94/2017 | [EPI_ISL_17767843](https://platform.epicov.org/epi3/start/EPI_ISL/17767843) | Russian Federation | 2017-Feb-28 | N.F. Gamaleya Research Center for Epidemiology and Microbiology | Import from public-domain | Voronina,O.L., Ryzhova,N.N., Aksenova,E.I., Kunda,M.S., Sharapova,N.A.N.E., Fedyakina,I.T., Chvala,I.A., Borisevich,S.V., Loguniv,D.Y.; Gintsburg,A.L. |
| 13 | A/wild duck/Poland/57/2017 | [EPI_ISL_300745](https://platform.epicov.org/epi3/start/EPI_ISL/300745) | Poland | 2017-Jan-27 |  | National Veterinary Research Institut Poland, PIWet-PIB | Swieton E., Smietanka K. |
| 14 | A/chicken/Greece/39_2017/2017 | [EPI_ISL_288362](https://platform.epicov.org/epi3/start/EPI_ISL/288362) | Greece | 2017-Feb-06 | Thessalonica Veterinary Centre (TVC) | Animal and Plant Health Agency (APHA) | Seekings, James; Ellis, Richard; Brookes, Sharon M; Reid, Stephen; Lewis, Nicola; Brown, Ian H; Dovas, C; Georgiades, D |
| 15 | A/turkey/England/003778/2017 | [EPI_ISL_253036](https://platform.epicov.org/epi3/start/EPI_ISL/253036) | United Kingdom | 2017-Jan-15 | Animal and Plant Health Agency (APHA) | Animal and Plant Health Agency (APHA) | Seekings, James; Ellis, Richard; Brookes, Sharon M; Reid, Scott; Essen, Stephen; Brown, Ian H |
| 16 | A/duck/Guizhou/S1321/2022(H5N1) | [EPI_ISL_12572656](https://platform.epicov.org/epi3/start/EPI_ISL/12572656) | China | 2022-Feb-22 | Harbin Veterinary Research Institute (CAAS) | Harbin Veterinary Research Institute (CAAS) | Pengfei Cui, Congcong Wang |
| 17 | A/Astrakhan/3212/2020 | [EPI_ISL_1038924](https://platform.epicov.org/epi3/start/EPI_ISL/1038924) | Russian Federation | 2020-Dec-12 | Center of Hygiene and Epidemiology in Astrakhan Region | State Research Center of Virology and Biotechnology (VECTOR) | Pyankova, O; Susloparov, I; Marchenko, V; Ryzhikov, A |
| 18 | A/goose/Czech Republic/18520-1/2021 | [EPI_ISL_17767177](https://platform.epicov.org/epi3/start/EPI_ISL/17767177) | Czech Republic | 2021-Sep-27 | State Veterinary Institute Prague | Import from public-domain | Nagy,A., Cernikova,L.; Stara,M. |
| 19 | A/chicken/Saitama/TU7-34,36/2021 | [EPI_ISL_15063425](https://platform.epicov.org/epi3/start/EPI_ISL/15063425) | Japan | 2021-Dec-07 |  | Import from public-domain | Soda,K.; Usui,T.; Ito,H.; Yamaguchi,T.; Ito,T. |
| 20 | A/teal/Miyazaki/211109-32/2021 | [EPI_ISL_15613494](https://platform.epicov.org/epi3/start/EPI_ISL/15613494) | Japan | 2021-Nov-09 |  | Import from public-domain | Soda,K.; Mekata,H.; Yamada,K.; Ito,H.; Usui,T.; Yamaguchi,T.; Ito,T. |
| 21 | A/common buzzard/Japan/2601B013/2022 | [EPI_ISL_16831015](https://platform.epicov.org/epi3/start/EPI_ISL/16831015) | Japan | 2022-Jan-27 |  | Import from public-domain | Soda,K.; Ito,H.; Usui,T.; Yamaguchi,T.; Ito,T. |
| 22 | A/quail/Korea/H526/2021 | [EPI_ISL_6959593](https://platform.epicov.org/epi3/start/EPI_ISL/6959593) | Korea, Republic of | 2021-Nov-08 | Animal and Plant Quarantine Agency (O-2144) | Animal and Plant Quarantine Agency (APQA) |  |
| 23 | A/peregrine falcon/Kanagawa/1409C001T1/2022 | [EPI_ISL_15923322](https://platform.epicov.org/epi3/start/EPI_ISL/15923322) | Japan | 2022-Sep-25 | National Institute for Environmental Studies | National Institute of Animal Health | Manabu,Onuma;Kei,Nabeshima;Atsushi,Haga;Hisako,Honjo;Misako,Yokoyama;Yuko,Uchida;Kohtaro,Miyazawa;Ryota,Tsunekuni;Junki,Mine;Saki,Sakuma;Asuka,Kumagai;Yoshihiro,Takadate |
| 24 | A/chicken/Ehime/TU11-2-24,25/2022 | [EPI_ISL_15063431](https://platform.epicov.org/epi3/start/EPI_ISL/15063431) | Japan | 2022-Jan-04 |  | Import from public-domain | Soda,K.; Ito,H.; Hisada,R.; Usui,T.; Yamaguchi,T.; Ito,T. |
| 25 | A/chicken/Kagoshima/21A6T/ 2021 | [EPI_ISL_6829533](https://platform.epicov.org/epi3/start/EPI_ISL/6829533) | Japan | 2021-Nov-12 | National Institute of Animal Health | National Institute of Animal Health |  |
| 26 | A/mandarin duck/Korea/WA585/2021 | [EPI_ISL_6959592](https://platform.epicov.org/epi3/start/EPI_ISL/6959592) | Korea, Republic of | 2021-Oct-26 | Animal and Plant Quarantine Agency (O-2144) | Animal and Plant Quarantine Agency (APQA) |  |
| 27 | A/duck/Guangdong/S4525/2021 | [EPI_ISL_12572655](https://platform.epicov.org/epi3/start/EPI_ISL/12572655) | China | 2021-Dec-08 | Harbin Veterinary Research Institute (CAAS) | Harbin Veterinary Research Institute (CAAS) | Pengfei Cui, Congcong Wang |
| 28 | A/duck/Hubei/SE220/2022 | [EPI_ISL_12572659](https://platform.epicov.org/epi3/start/EPI_ISL/12572659) | China | 2022-Jan-10 | Harbin Veterinary Research Institute (CAAS) | Harbin Veterinary Research Institute (CAAS) | Pengfei Cui, Congcong Wang |
| 29 | A/duck/Hulu Sungai Utara/A0522064-06/2022 | [EPI_ISL_17371282](https://platform.epicov.org/epi3/start/EPI_ISL/17371282) | Indonesia | 2022-Apr-04 | Disease Investigation Centre Regional V Banjarbaru (BPPVRV) | Balai Besar Veteriner Wates | Wibawa, Hendra; Wibowo, Putut Eko; Lestari, -; Irianingsih, Sri Handayani; Supriyadi, Arif; Fiqri, Anna Januar; Fahmia, Zaza; Silaban, Jesiaman; Mulyawan, Herdiyanto |
| 30 | A/duck/Hulu Sungai Utara/A0522064-03-04/2022 | [EPI_ISL_17371283](https://platform.epicov.org/epi3/start/EPI_ISL/17371283) | Indonesia | 2022-Apr-04 | Disease Investigation Centre Regional V Banjarbaru (BPPVRV) | Balai Besar Veteriner Wates | Wibawa, Hendra; Wibowo, Putut Eko; Lestari, -; Irianingsih, Sri Handayani; Supriyadi, Arif; Fiqri, Anna Januar; Fahmia, Zaza; Silaban, Jesiaman; Mulyawan, Herdiyanto |
| 31 | A/duck/Hulu Sungai Utara/A0522067-06-07/2022 | [EPI_ISL_17371284](https://platform.epicov.org/epi3/start/EPI_ISL/17371284) | Indonesia | 2022-Apr-04 | Disease Investigation Centre Regional V Banjarbaru (BPPVRV) | Balai Besar Veteriner Wates | Wibawa, Hendra; Wibowo, Putut Eko; Lestari, -; Irianingsih, Sri Handayani; Supriyadi, Arif; Fiqri, Anna Januar; Fahmia, Zaza; Silaban, Jesiaman; Mulyawan, Herdiyanto |
| 32 | A/domestic duck/Hungary/7341/2015 | [EPI_ISL_177584](https://platform.epicov.org/epi3/start/EPI_ISL/177584) | Hungary | 2015-Feb-23 | Danam.Vet.Molbiol | Danam.Vet.Molbiol | Krisztian,Banyai; Szilvia,Farkas; Adam,Dan |
| 33 | A/chicken/Netherlands/14015766/2014 | [EPI_ISL_174349](https://platform.epicov.org/epi3/start/EPI_ISL/174349) | Netherlands | 2014-Nov-19 | Wageningen Bioveterinary Research | Wageningen Bioveterinary Research | Heutink, Rene; Harders, Frank; Verschuren-Pritz, Sylvia; Bossers, Alex; Koch, Guus; Bouwstra, Ruth |
| 34 | A/wild bird/Korea/H2291/2015 | [EPI_ISL_234336](https://platform.epicov.org/epi3/start/EPI_ISL/234336) | Korea, Republic of | 2015-Jan-30 |  | Animal and Plant Quarantine Agency (APQA) |  |
| 35 | A/mallard/Oregon/195536/2014 | [EPI_ISL_206441](https://platform.epicov.org/epi3/start/EPI_ISL/206441) | United States | 2014-Dec-24 |  | Import from public-domain | Killian,M.L.; Ip,H.S.; Griffin,K.; Messer,J.; McMullen,K.; Dusek,R.; Bodenstein,B. |
| 36 | A/mallard/Idaho/AH0007413/2015 | [EPI_ISL_206408](https://platform.epicov.org/epi3/start/EPI_ISL/206408) | United States | 2015-Jan-17 |  | Import from public-domain | Killian,M.L. |
| 37 | A/Canada goose/Kansas/197850/2015 | [EPI_ISL_206450](https://platform.epicov.org/epi3/start/EPI_ISL/206450) | United States | 2015-Mar-13 |  | Import from public-domain | Killian,M.L.; Ip,H.S.; Griffin,K.; Messer,J.; McMullen,K.; Long,R.; Hesting,S. |
| 38 | A/turkey/Iowa/15-013179-4/2015 | [EPI_ISL_301110](https://platform.epicov.org/epi3/start/EPI_ISL/301110) | United States | 2015-Jan-01 |  | Import from public-domain | Lee,D.-H.; Torchetti,M.; Hicks,J.; Killian,M.; Bahl,J.; Pantin-Jackwood,M.; Swayne,D. |
| 39 | A/Environment/Shenzhen/1/2015 | [EPI_ISL_205314](https://platform.epicov.org/epi3/start/EPI_ISL/205314) | China | 2015-Dec-21 |  | WHO Chinese National Influenza Center | Fang, Shisong;Yang, Lei |
| 40 | A/Shenzhen/1/2015 | [EPI_ISL_205313](https://platform.epicov.org/epi3/start/EPI_ISL/205313) | China | 2015-Dec-28 | Shenzhen center for disease control and prevention | WHO Chinese National Influenza Center | Fang, Shisong;Yang, Lei |
| 41 | A/Environment/Guangdong/40113/2015 | [EPI_ISL_219808](https://platform.epicov.org/epi3/start/EPI_ISL/219808) | China | 2015-May-11 |  | WHO Chinese National Influenza Center | Wang,Dayan;Li,Xiaodan;Zou,Shumei;Zhang,Ye;Bo,Hong;Li,Xiyan;Chen,Wenbing;Yang,Lei;Shu,Yuelong |
| 42 | A/poultry/China/XY165.4/2016 | [EPI_ISL_17767176](https://platform.epicov.org/epi3/start/EPI_ISL/17767176) | China | 2016-Sep-01 | Chinese Academy of Medical Sciences | Import from public-domain | Zhao,Z. |
| 43 | A/chicken/Guangdong/GD1602/2016 | [EPI_ISL_282397](https://platform.epicov.org/epi3/start/EPI_ISL/282397) | China | 2016-Mar-22 |  | Import from public-domain | Sun,W. |
| 44 | A/chicken/Anhui/MZ33/2016 | [EPI_ISL_297930](https://platform.epicov.org/epi3/start/EPI_ISL/297930) | China | 2016-Feb-01 |  | Import from public-domain | Liu,K.; Gu,M.; Gao,R.; Li,J.; Liu,D.; Sun,W.; Hu,J.; Xu,X.; Wang,X.; Liu,X. |
| 45 | A/black swan/Akita/1/2016 | [EPI_ISL_243058](https://platform.epicov.org/epi3/start/EPI_ISL/243058) | Japan | 2016-Nov-19 |  | Import from public-domain | Okamatsu,M.; Hiono,T.; Matsuno,K.; Kida,H.; Sakoda,Y. |
| 46 | A/whooper swan/Iwate/5/2016 | [EPI_ISL_17767791](https://platform.epicov.org/epi3/start/EPI_ISL/17767791) | Japan | 2016-Dec-18 | Graduate School of Veterinary Medicine, Hokkaido University | Import from public-domain | Sakoda,Y., Okamatsu,M.; Matsuno,K. |
| 47 | A/whooper swan/Niigata/13/2017 | [EPI_ISL_17767801](https://platform.epicov.org/epi3/start/EPI_ISL/17767801) | Japan | 2017-Jan-27 | Graduate School of Veterinary Medicine, Hokkaido University | Import from public-domain | Sakoda,Y., Okamatsu,M.; Matsuno,K. |
| 48 | A/coot/Iwate/13/2016 | [EPI_ISL_256512](https://platform.epicov.org/epi3/start/EPI_ISL/256512) | Japan | 2016-Dec-22 |  | Hokkaido University |  |
| 49 | A/chicken/Miyazaki/2-3C/2017 | [EPI_ISL_243687](https://platform.epicov.org/epi3/start/EPI_ISL/243687) | Japan | 2017-Jan-24 |  | National Institute of Animal Health |  |
| 50 | A/chicken/Korea/HN1/2016(H5N6) | [EPI_ISL_239261](https://platform.epicov.org/epi3/start/EPI_ISL/239261) | Korea, Republic of | 2016-Nov-16 |  | Animal and Plant Quarantine Agency (APQA) |  |
| 51 | A/duck/Korea/ES2/2016(H5N6) | [EPI_ISL_239262](https://platform.epicov.org/epi3/start/EPI_ISL/239262) | Korea, Republic of | 2016-Nov-16 |  | Animal and Plant Quarantine Agency (APQA) |  |
| 52 | A/chicken/Kumamoto/1-5T/2016 | [EPI_ISL_241779](https://platform.epicov.org/epi3/start/EPI_ISL/241779) | Japan | 2016-Dec-27 |  | National Institute of Animal Health | Saito, T.; Takemae, N. |
| 53 | A/turtledove/Wuhan/HKBJ43/2015 | [EPI_ISL_205140](https://platform.epicov.org/epi3/start/EPI_ISL/205140) | China | 2015-Jan-01 |  | Import from public-domain | Chen,L.-J.; Lin,X.-D.; Guo,W.-P.; Tian,J.-H.; Zhang,Y.-Z. |
| 54 | A/duck/Wuhan/WHYF02/2015 | [EPI_ISL_205115](https://platform.epicov.org/epi3/start/EPI_ISL/205115) | China | 2015-Jan-01 |  | Import from public-domain | Chen,L.-J.; Lin,X.-D.; Guo,W.-P.; Tian,J.-H.; Zhang,Y.-Z. |
| 55 | A/Environment/Guangdong/33311/2015 | [EPI_ISL_219813](https://platform.epicov.org/epi3/start/EPI_ISL/219813) | China | 2015-Mar-24 |  | WHO Chinese National Influenza Center | Wang,Dayan;Li,Xiaodan;Zou,Shumei;Zhang,Ye;Bo,Hong;Li,Xiyan;Chen,Wenbing;Yang,Lei;Shu,Yuelong |
| 56 | A/duck/Hunan/01.21 YYFQH006-O/2015(H5N6) | [EPI_ISL_199079](https://platform.epicov.org/epi3/start/EPI_ISL/199079) | China | 2015-Jan-21 |  | Institute of Microbiology, Chinese Academy of Sciences |  |
| 57 | A/goose/Anhui/AH350/2015 | [EPI_ISL_255822](https://platform.epicov.org/epi3/start/EPI_ISL/255822) | China | 2015-Dec-19 |  | Wuhan Institute of Virology |  |
| 58 | A/muscovy duck/Vietnam/LBM810/2015 | [EPI_ISL_230756](https://platform.epicov.org/epi3/start/EPI_ISL/230756) | Vietnam | 2015-Nov-10 |  | Import from public-domain | Soda,K.; Nguyen,K.H.; Le,Q.M.; Ito,T.; Maegaki,S. |
| 59 | A/chicken/Vietnam/HU4-42/2015 | [EPI_ISL_293989](https://platform.epicov.org/epi3/start/EPI_ISL/293989) | Vietnam | 2015-Nov-07 |  | Import from public-domain | Sakoda,Y.; Okamatsu,M.; Matsuno,K.; Jizou,M. |
| 60 | A/Environment/Hunan/07767/2015 | [EPI_ISL_219816](https://platform.epicov.org/epi3/start/EPI_ISL/219816) | China | 2015-Jan-08 |  | WHO Chinese National Influenza Center | Wang,Dayan;Li,Xiaodan;Zou,Shumei;Zhang,Ye;Bo,Hong;Li,Xiyan;Chen,Wenbing;Yang,Lei;Shu,Yuelong |
| 61 | A/duck/Hunan/HN42/2015 | [EPI_ISL_255764](https://platform.epicov.org/epi3/start/EPI_ISL/255764) | China | 2015-Apr-06 |  | Wuhan Institute of Virology |  |
| 62 | A/goose/Hunan/110/2014 | [EPI_ISL_255827](https://platform.epicov.org/epi3/start/EPI_ISL/255827) | China | 2014-Nov-13 |  | Wuhan Institute of Virology |  |
| 63 | A/duck/Eastern China/S0908/2014 | [EPI_ISL_208838](https://platform.epicov.org/epi3/start/EPI_ISL/208838) | China | 2014-Sep-08 |  | Import from public-domain | Sun,H.; Sun,Y.; Pu,J.; Liu,L.; Li,C.; Xu,G.; Qin,M.; Zhang,Y.; Zhao,H.; Wei,K.; Liu,J. |
| 64 | A/duck/Hunan/233/2014 | [EPI_ISL_255493](https://platform.epicov.org/epi3/start/EPI_ISL/255493) | China | 2014-Nov-13 |  | Wuhan Institute of Virology |  |
| 65 | A/duck/Hunan/HN13/2015 | [EPI_ISL_255529](https://platform.epicov.org/epi3/start/EPI_ISL/255529) | China | 2015-Apr-06 |  | Wuhan Institute of Virology |  |
| 66 | A/duck/Hunan/HN334/2015 | [EPI_ISL_255757](https://platform.epicov.org/epi3/start/EPI_ISL/255757) | China | 2015-Dec-18 |  | Wuhan Institute of Virology |  |
| 67 | A/duck/Hunan/HN336/2015 | [EPI_ISL_255759](https://platform.epicov.org/epi3/start/EPI_ISL/255759) | China | 2015-Dec-18 |  | Wuhan Institute of Virology |  |

We gratefully acknowledge the authors, originating and submitting laboratories of the sequences from GISAID’s EpiFlu™ Database on which this research is based.
